# Differential behavioral effects of systemic oxytocin receptor blockade across social tasks and hierarchical ranks in male mice

**DOI:** 10.1101/2022.07.23.501222

**Authors:** Daiki Nasukawa, Shuntaro Matsushima, Kota Yamada, Yusuke Ujihara, Haruka Hirakata, Ryuto Tamura, Saya Yatagai, Kazuko Hayashi, Koji Toda

## Abstract

Oxytocin receptor signaling has been implicated in diverse social behaviors, but whether its behavioral effects depend on social tasks and an individual’s position within a hierarchy remains unclear. We examined the effects of systemic administration of L-368,899, a blood-brain barrier-penetrating oxytocin receptor antagonist, across four domains of social behavior in male C57BL/6J mice: social rank, sex preference, general social preference, and dyadic interaction. Following the establishment of stable hierarchies in the tube test, administration of L-368,899 (10 mg/kg, intraperitoneally) to first-ranked mice did not alter their hierarchical position. Rank fluctuations were observed more frequently when the antagonist was administered to second-ranked mice. In both rank conditions, the remaining cage mates received saline. In the sex-preference task, saline-treated males spent more time in the female stimulus zone than in the male stimulus zone, whereas this preference was not statistically detectable following treatment with L-368,899 (3 or 10 mg/kg). In contrast, L-368,899 did not significantly alter proximity-based preference for a novel male over a nonsocial object or affect contact duration and inter-individual distance during dyadic interactions between familiar male cage mates, in which both members of each pair received the same treatment. Thus, systemic oxytocin receptor blockade did not produce a generalized reduction in social approach. Instead, the observed behavioral patterns suggest that its effects depend on specific social behavioral tasks and hierarchical position. These findings provide a pharmacological basis for further investigation of the task- and rank-dependent contributions of oxytocin receptor signaling to social behavior.

## 1. Introduction

Oxytocin is an important neuromodulator of social behavior (Donaldson and Young, 2008) and has been implicated in pair bonding (Young and Wang, 2004), maternal behavior (Insel, 1990), aggression (Bosch et al., 2005), social decision-making (Chang et al., 2012), observational fear conditioning (Pisansky et al., 2017), and social recognition (Ferguson et al., 2000). Accordingly, oxytocin signaling has been investigated in relation to neuropsychiatric disorders characterized by social dysfunction, including autism spectrum disorder (Andari et al., 2020), social anxiety and social fear (Neumann and Slattery, 2016), and schizophrenia (Shilling and Feifel, 2016). These diverse associations raise the question of whether oxytocin receptor signaling contributes similarly to different forms of social behavior or whether its contribution varies with the behavioral demands and social relationships involved. Examining responses to the same pharmacological manipulation across several behavioral tasks provides a way to investigate this question, while recognizing that such comparisons involve differences in both social context and the behaviors measured.

Social rank emerges through repeated interactions among individuals living within a group. Establishing a stable social hierarchy can reduce unnecessary conflict and the energetic costs of competition while determining access to limited resources and living spaces. Consequently, social rank can influence survival, reproductive success, health, and a wide range of behaviors (Wang et al., 2014). Oxytocin receptor densities in several brain regions differ among mice occupying different positions within a social hierarchy (Lee et al., 2019), raising the possibility that responses to oxytocin receptor blockade also vary with social rank. However, it remains unclear whether acute pharmacological blockade of these receptors has different effects on established hierarchical relationships depending on the rank of the treated individual.

The tube test is a well-established behavioral assay for evaluating social hierarchy in rodents (Fan et al., 2019; Wang et al., 2014). In this test, two mice enter opposite ends of a narrow tube, and the outcome is determined when one mouse retreats or is pushed out by the other. Hierarchical relationships are determined from the outcomes of round-robin competitions among cage mates. Social rank measured using the tube test is associated with several ethologically relevant behaviors. Dominant mice show increased allogrooming and whisker trimming of subordinate animals (Wang et al., 2011), produce more courtship ultrasonic vocalizations toward females than subordinate males (Nyby et al., 1976; Wang et al., 2011), and spend more time occupying a limited warm spot in the hot-spot competition task (Zhou et al., 2017). These associations support the use of the tube test to assess established social hierarchies and to track the rank trajectories of treated individuals and their cage mates following pharmacological manipulation.

To examine whether responses to oxytocin receptor blockade extend beyond competitive interactions, complementary tasks are needed to assess other forms of social behavior. A sex-preference task compares the time a male subject spends near female and male conspecifics, providing a measure of proximity-based preference between the sexes. A social-preference task compares time spent near an unfamiliar male conspecific and a nonsocial object, allowing assessment of social approach without an opposite-sex stimulus. These two tasks use similar spatial measures but differ in the choice presented to the subject. Neither measure alone establishes the animal’s motivation or the amount of active stimulus investigation. A dyadic interaction task provides a complementary assessment by allowing familiar male cage mates to interact freely and quantifying their contact duration and inter-individual distance. In combination with the tube test, these tasks permit examination of several components of social behavior while retaining the distinction between competitive outcomes, proximity-based preferences, and direct interactions.

In the present study, we used L-368,899, a nonpeptide oxytocin receptor antagonist, to examine behavioral responses to systemic oxytocin receptor blockade across these four tasks in male mice. L-368,899 exhibits high affinity for rat and human uterine oxytocin receptors (Ki = 3.6 and 13 nM, respectively), with selectivity over vasopressin V1 and V2 receptors in tissue-based binding assays from both species (Pettibone et al., 1993). These pharmacological properties support its use as an oxytocin receptor antagonist, although they do not establish in vivo receptor selectivity under the conditions used in the present mouse experiments. We examined established social rank, proximity-based sex preference, general social preference, and dyadic interactions between familiar male cage mates. In the tube test, L-368,899 was administered to either the first- or second-ranked mouse, while the remaining cage mates received saline. In the preference tasks, only subject mice received saline or L-368,899, and stimulus mice received no injections. In the dyadic interaction task, both members of each pair received the same treatment. This design allowed us to examine whether behavioral responses to L-368,899 varied across social tasks and according to the hierarchical position of the treated individual.

## 2. Materials and methods

### 2.1. Animals

All data were collected from 2–6-month-old adult male wild-type C57BL/6J mice. The mice were maintained on a 12:12 light cycle. All the experiments were conducted during the dark phase of the light cycle. The mice had unrestricted access to food and water in their cages. Their weights were monitored daily. All animal procedures were approved by the Animal Research Committee of Keio University. This study is reported in accordance with ARRIVE guidelines (https://arriveguidelines.org).

### 2.2. Experimental procedures

#### 2.2.1. Experiment 1: Tube test

In Experiment 1, we examined the effects of intraperitoneal administration of the oxytocin receptor antagonist L-368,899 on established social rank. Thirty-two adult male C57BL/6J mice, aged 3–6 months, were housed in eight cages, with four mice per cage. The same eight cages were tested sequentially under both treatment conditions: administration of L-368,899 to the first-ranked mouse and administration to the second-ranked mouse. In all cages, the first-ranked-mouse condition was conducted first, followed by the second-ranked-mouse condition after an interval of at least 1 week. Before each treatment condition, the hierarchy was reassessed, and L-368,899 was administered only after the ranks of all four mice had remained stable for three consecutive days. The tube-test procedure was based on the protocol described by Fan et al. (2019), with some modifications. The experiment consisted of habituation, training, and testing phases. During habituation, a short acrylic tube (35 mm inner diameter, 2 mm wall thickness, and 10 cm long; Figure 1A) was placed in the home cage for 2 h per day on two consecutive days, allowing the mice to explore it freely (Figure 1B; Supplementary Movie 1). During training, a longer acrylic tube (35 mm inner diameter, 2 mm wall thickness, and 30 cm long; Figure 1A) was used. Each mouse was trained to pass through the tube 10 times per day, five times from each entrance, on two consecutive days (Figure 1C; Supplementary Movie 2). During testing, all possible pairs of cage mates competed in a round-robin design (Figure 1D). Because the apparatus had left and right starting positions, each pair competed twice, once from each side. The order of matches was counterbalanced across testing sessions and randomized within the counterbalancing scheme using a random-number generator. Testing was conducted in a chamber measuring 63.5 × 40.0 × 60.0 cm (MED Associates), with the chamber door kept open. White noise (75 dB) was presented throughout the experiment to mask external noise, and behavior was recorded using cameras positioned 30 cm from the tube (Logicool HD Webcam C920r, Logicool Co., Ltd., Tokyo, Japan). At the beginning of each match, the experimenter simultaneously released the two mice from opposite entrances by releasing their tails (Supplementary Movie 3). A mouse was classified as the loser when it retreated from the tube or was pushed backward until one of its hind limbs exited the tube; the other mouse was classified as the winner. No time limit was imposed. If each mouse won one of the two matches against the same opponent, the pairwise outcome was classified as a draw. Social rank was determined from the total number of wins in the round-robin competition. A hierarchy was considered stable when the ranks of all four mice remained unchanged for three consecutive testing days. The order of the mice within each hierarchy was classified as first, second, third, or fourth rank. Once the hierarchy had stabilized, all four mice received an intraperitoneal injection of saline 30 min before each baseline testing session to control for the effects of handling, needle penetration, and injection. On the day following confirmation of rank stability, the target-ranked mouse received L-368,899 at 10 mg/kg, whereas the other three cage mates received saline. Each mouse was returned to its home cage immediately after injection, and the tube test was conducted 30 min later. Testing continued for 3 days after antagonist administration, with all four mice receiving saline on the subsequent testing days. In every cage, L-368,899 was first administered to the first-ranked mouse. After completion of this condition, an interval of at least 1 week was allowed before the second-ranked-mouse condition. During this interval, the hierarchy was reassessed, and the second-ranked-mouse condition was initiated only after all four ranks had again remained stable for three consecutive days. The same treatment and testing procedures were then repeated, except that L-368,899 was administered to the second-ranked mouse. Thus, all eight cages completed both treatment conditions in a fixed order, with the first-ranked-mouse condition preceding the second-ranked-mouse condition. The number of wins and losses and the duration of each match were recorded manually by an observer blinded to treatment condition. Body weights were measured throughout the experiment.

**Figure 1.**
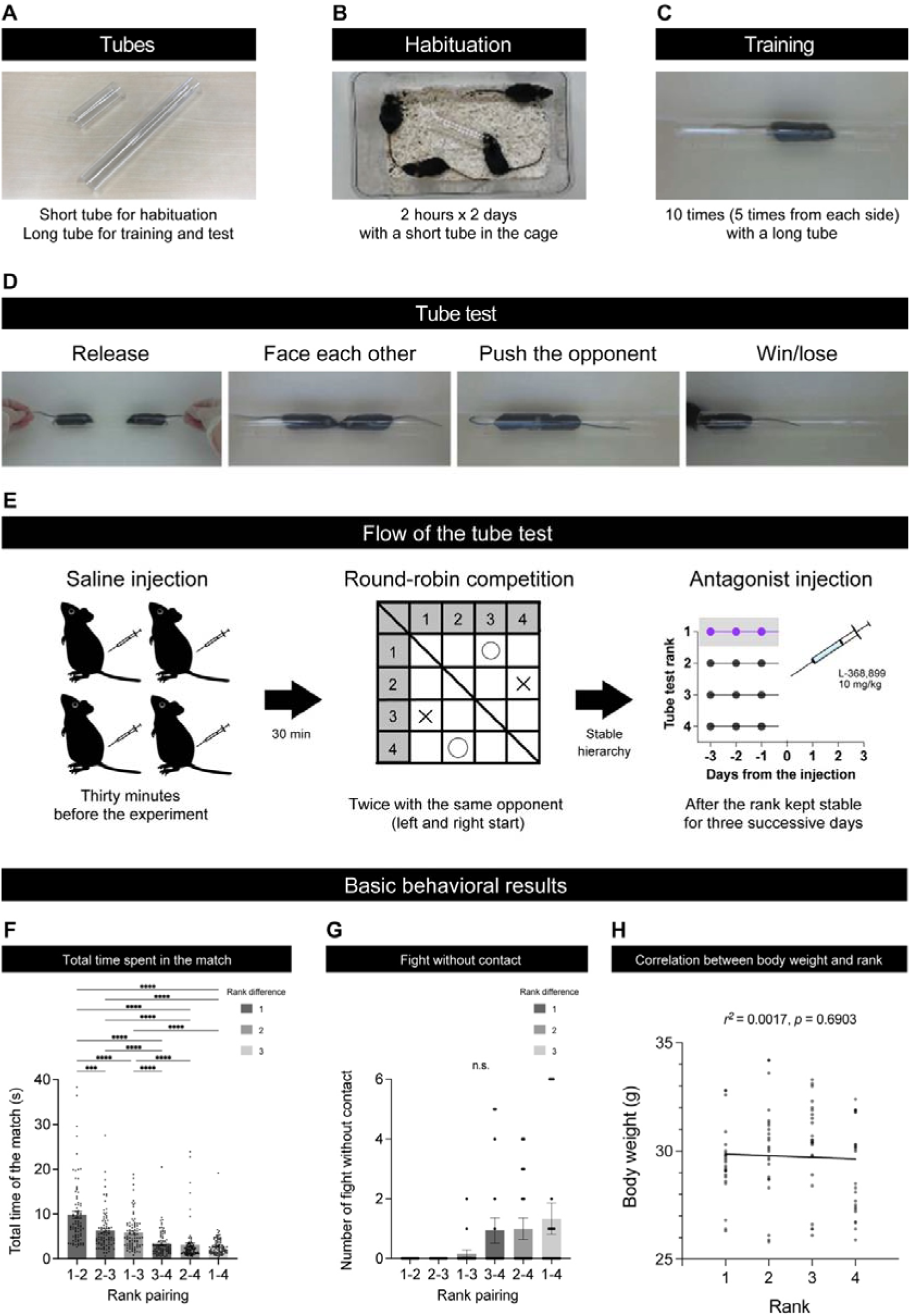
Tube-test procedure and pretreatment behavioral characteristics. A. Acrylic tubes used for habituation and testing. A short tube (35 mm inner diameter, 2 mm wall thickness, and 10 cm long) was used during habituation, and a longer tube (35 mm inner diameter, 2 mm wall thickness, and 30 cm long) was used during training and testing. B. Habituation procedure. A short tube was placed in each home cage for 2 h per day on two consecutive days, allowing the four cage mates to explore it freely. C. Training procedure. Each mouse was trained to pass through the long tube 10 times per day, five times from each entrance, on two consecutive days. D. Tube-test procedure. Two cage mates were released simultaneously from opposite ends of the tube. The mouse that retreated or was pushed backward until one of its hind limbs exited the tube was classified as the loser, and the other mouse was classified as the winner. Each pair competed twice, once from each starting side. A pairwise outcome was classified as a draw when each mouse won one match. E. Experimental timeline. All possible pairs of cage mates competed in a round-robin design. A hierarchy was considered stable when the ranks of all four mice remained unchanged for three consecutive days. All four mice received saline 30 min before baseline testing. On the administration day, the target-ranked mouse received L-368,899 (10 mg/kg, intraperitoneally), whereas the other three cage mates received saline. Testing began 30 min after injection. All eight cages completed the first-ranked-mouse condition first, followed by the second-ranked-mouse condition after an interval of at least 1 week and confirmation of stable ranks for three consecutive days. F. Match duration for each rank pairing before antagonist administration. Match duration was measured from the simultaneous release of the mice until the outcome was determined using the criteria described in D. Each point represents an individual match. G. Number of tube-test matches that ended without direct physical contact between the two mice, shown separately for each rank pairing during the pretreatment period. H. Relationship between body weight and social rank before antagonist administration. The plotted dataset contains 96 measurements from 32 mice, and the line represents the fitted simple linear regression. Panels F–H describe pretreatment characteristics and do not represent comparisons before and after L-368,899 administration. Post-administration rank trajectories are shown in Figure 2. Bars and error bars represent the mean ± SEM where shown. \*\*\*\**p* < 0.0001, \*\*\**p* < 0.001.

**Figure 2.**
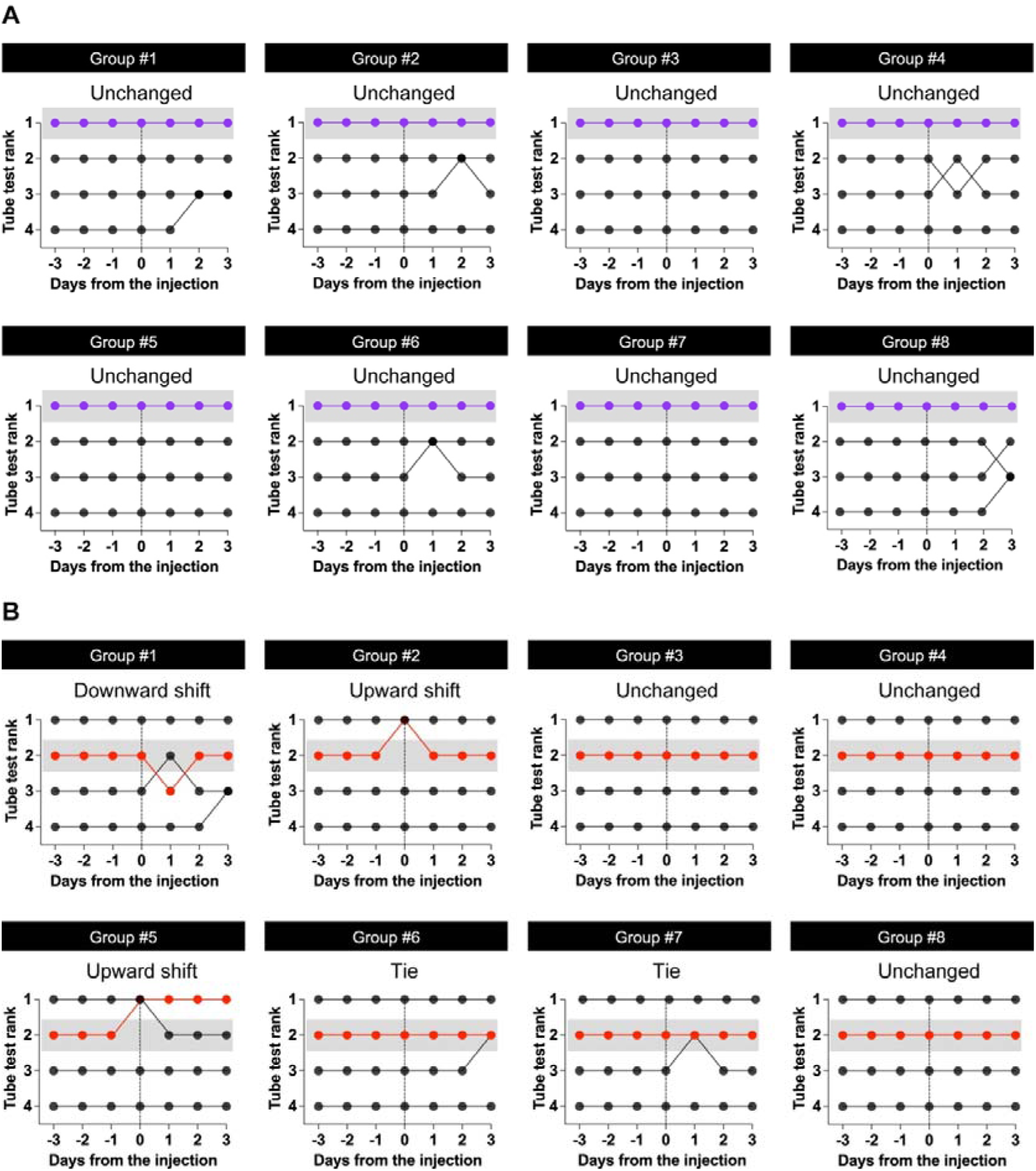
Individual rank trajectories following L-368,899 administration to first- or second-ranked mice. Tube-test ranks are shown for three days before administration, the administration day (Day 0), and three subsequent days. On Day 0, the target-ranked mouse received L-368,899 (10 mg/kg, intraperitoneally), whereas the other three cage mates received saline. All four mice received saline on the baseline and subsequent testing days. Testing began 30 min after injection. A. Administration to first-ranked mice. The treated mouse retained first rank without becoming tied with another cage mate in all eight cages, although rank changes occurred among some untreated cage mates. B. Administration to second-ranked mice. Upward shifts occurred on Day 0 in Groups #2 and #5. The treated mouse returned to second rank on Day 1 in Group #2 but remained at first rank throughout the remaining observation period in Group #5. In Group #1, the treated mouse fell to third rank on Day 1 and returned to second rank on Day 2. In Groups #6 and #7, the treated mouse became tied with the initially third-ranked cage mate on Day 3 and Day 1, respectively. In Groups #3, #4, and #8, the treated mouse remained at second rank without becoming tied with another cage mate. Colored lines identify the L-368,899-treated mice, gray shading marks their pretreatment rank, and vertical dashed lines indicate Day 0. Labels above each panel describe the outcome of the treated mouse during Days 0–3, rather than changes involving only untreated cage mates. Each pair of mice competed twice per session; an outcome of one win and one loss was classified as a draw. Tied ranks are displayed at the same vertical position. The same eight cages completed both conditions, with treatment of first-ranked mice preceding treatment of second-ranked mice. A change in the treated mouse’s numerical rank or the occurrence of a tie was observed in 0/8 cages in A and 5/8 cages in B (two-sided exact McNemar test, *p* = 0.0625).

#### 2.2.2. Experiment 2: Sex preference task

To examine whether oxytocin receptor antagonists affect sex preference in male mice, we used a place preference task (Figure 3). Twenty-four adult male C57BL/6J mice aged 2–4 months were used as subjects, and six adult male and female C57BL/6J mice aged 2–4 months were used as stimuli. All subject mice were group-housed before testing; none were individually housed. A custom-made white vinyl chloride box (25 cm length × 50 cm width × 30 cm height) was used for the test. Stimulus mice were placed in white cylindrical stimulus baskets (8 cm diameter × 10 cm height), which were positioned at the center of opposite ends of the 50-cm-wide box. A 500-mL empty glass bottle was placed on top of each basket to prevent the subject mouse from climbing. Two partition walls (5 cm length × 5 mm thickness × 30 cm height) were attached to both sides of the center of the 50-cm-wide box to partially separate the left and right spaces. A camera (Logicool HD Pro Webcam C920r, Logicool Co., Ltd., Tokyo, Japan) was positioned 80 cm above the floor of the box, and behavior was recorded using a Windows PC. Subject mice received an intraperitoneal injection of saline or L-368,899 (3 or 10 mg/kg). The male and female stimulus mice received no injections. Thirty minutes after injection, each subject mouse was placed in the apparatus and allowed to habituate for 15 min. The subject mouse was then temporarily removed, and the male and female stimulus mice were placed in their respective stimulus baskets. The subject mouse was returned to the center of the apparatus and allowed to explore freely for 10 min. White noise (75 dB) was presented throughout the experiment to mask external noise.

**Figure 3.**
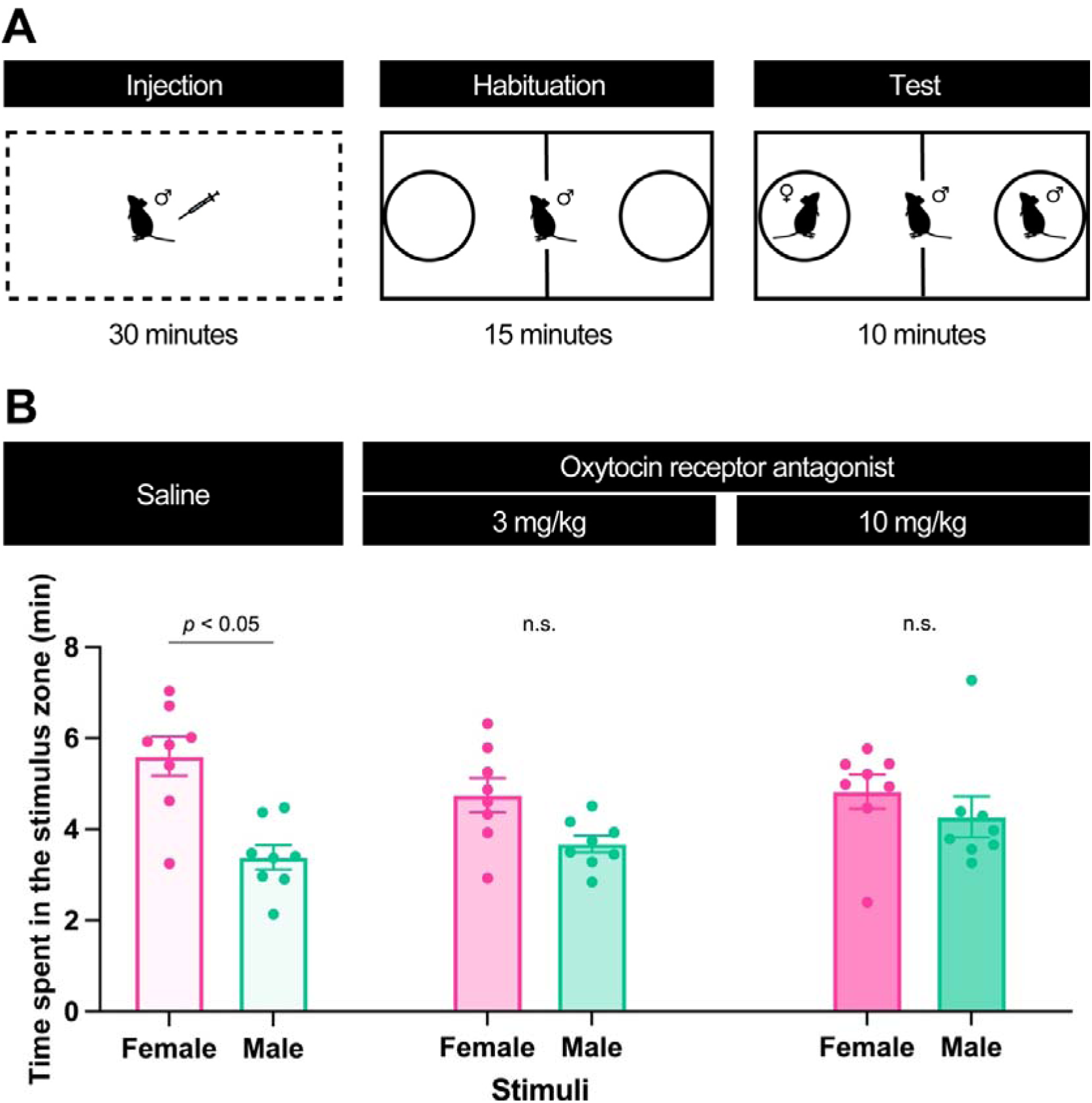
Effect of the oxytocin receptor antagonist on the sex preference task. A. Schematic of the sex-preference test. The apparatus consisted of a custom-made white vinyl chloride box (25 × 50 × 30 cm) divided into two compartments by a central wall, with a wire basket containing a stimulus mouse positioned at each end. Only subject mice received an intraperitoneal injection of saline or L-368,899 (3 or 10 mg/kg); the male and female stimulus mice received no injections. Subject mice were returned to their home cage after injection. Thirty minutes after injection, each subject mouse was allowed to habituate to the apparatus for 15 min. A male and a female stimulus mouse were then placed in the baskets, and the subject mouse was allowed to explore freely for 10 min. White noise (75 dB) was presented throughout the experiment. B. Time spent in the male and female stimulus zones. Zone occupancy was defined by the body centroid being located within the predefined 7.5-cm-wide zone adjacent to each stimulus basket, without requiring stimulus-directed orientation or active sniffing. Saline-treated mice spent significantly more time in the female stimulus zone than in the male stimulus zone. In contrast, a statistically significant preference for the female stimulus zone was not detected following treatment with either 3 or 10 mg/kg L-368,899. However, the drug × stimulus interaction was not statistically significant. Data are presented as the mean ± SEM. Saline: *n* = 8; L-368,899 3 mg/kg: *n* = 8; L-368,899 10 mg/kg: *n* = 8. Statistical significance was assessed using two-way mixed-design ANOVA followed by Šidák’s multiple-comparisons test.

#### 2.2.3. Experiment 3: Social preference task

To examine whether systemic oxytocin receptor blockade alters general social preference, we used a social-preference task in which subject mice could investigate an unfamiliar male conspecific or a novel nonsocial object (Figure 4; Tamura et al., 2026; Inoue et al., 2026). Eighteen adult male C57BL/6J mice, aged 2–4 months, were used as subjects and assigned to either the saline-treated group (*n* = 9) or the L-368,899-treated group (*n* = 9). Two adult male C57BL/6J mice, aged 3 months, were used as social stimuli. All subject mice were group-housed before testing; none were individually housed. The apparatus consisted of a custom-made white vinyl chloride box measuring 25 × 50 × 30 cm. Two white cylindrical stimulus baskets (8 cm in diameter and 10 cm high) were positioned at the centers of the opposite ends of the 50-cm-long apparatus. Each basket contained a male or female stimulus mouse. A 500-mL empty glass bottle was placed on top of each stimulus basket to prevent the subject mouse from climbing onto it. Two partition walls, each measuring 5 cm long × 5 mm thick × 30 cm high, were attached near the center of the apparatus to partially separate the left and right stimulus areas. A camera (Logicool HD Pro Webcam C920r; Logicool Co., Ltd., Tokyo, Japan) was positioned 80 cm above the floor of the apparatus, and behavior was recorded using a Windows PC. Subject mice received an intraperitoneal injection of saline or L-368,899 (10 mg/kg). The male stimulus mice received no injections. Thirty minutes after injection, each subject mouse was placed in the apparatus and allowed to habituate for 15 min. The subject mouse was then temporarily removed, and an unfamiliar male C57BL/6J mouse and a novel nonsocial object were placed in the respective stimulus baskets. The subject mouse was returned to the center of the apparatus and allowed to explore freely for 10 min. White noise (75 dB) was presented throughout the experiment to mask external noise.

**Figure 4.**
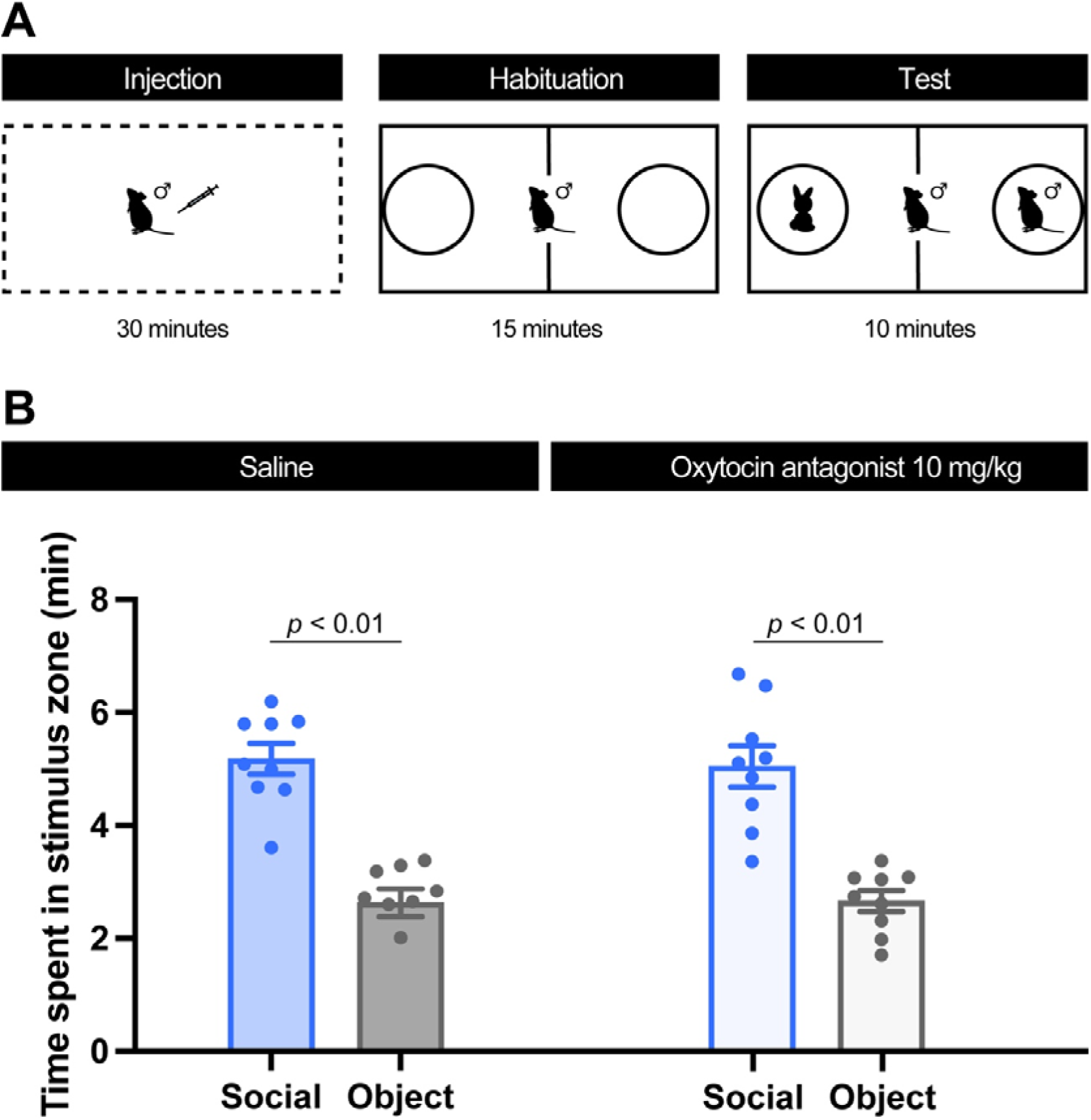
Effect of the oxytocin receptor antagonist on the social preference task. A. Schematic of the social-preference test. The apparatus consisted of a custom-made white vinyl chloride box (25 × 50 × 30 cm) divided into two compartments by a central wall, with a stimulus basket positioned at each end. Only subject mice received an intraperitoneal injection of saline or L-368,899 (10 mg/kg); the male stimulus mice received no injections. Thirty minutes after injection, the subject mice were allowed to habituate to the apparatus for 15 min. An unfamiliar male mouse and a novel nonsocial object were then placed in the respective baskets, and the subject mouse was allowed to explore freely for 10 min. White noise (75 dB) was presented throughout the experiment. B. Time spent in the social and nonsocial stimulus zones. Zone occupancy was defined by the body centroid being located within the predefined 7.5-cm-wide zone adjacent to each stimulus basket, without requiring stimulus-directed orientation or active sniffing. Mice spent significantly more time in the zone adjacent to the unfamiliar male than in the zone adjacent to the nonsocial object in both treatment groups. Neither the main effect of drug nor the stimulus × drug interaction was statistically significant. Data are presented as the mean ± SEM. Saline: *n* = 9; L-368,899: *n* = 9. Statistical significance was assessed using two-way mixed-design ANOVA followed by Šidák’s multiple-comparisons test.

#### 2.2.4. Experiment 4: Dyadic interaction in the open-field box

To examine whether systemic oxytocin receptor blockade alters dyadic social interactions between familiar male cage mates, we observed pairs of mice in an open-field arena using a previously described procedure (Figure 5; Supplementary Movie 4; Yamamoto et al., 2026). Twenty adult male C57BL/6J mice, aged 2–3 months, were housed in pairs of familiar cage mates. The resulting 10 pairs were assigned to either the saline-treated condition (n = 5 pairs; 10 mice) or the L-368,899-treated condition (n = 5 pairs; 10 mice). The apparatus consisted of a custom-made white vinyl chloride open-field arena measuring 50 × 50 × 50 cm. A camera (HD Webcam C270n; Logicool Co., Ltd., Tokyo, Japan) was positioned 106 cm above the floor of the arena, and behavior was recorded using a Windows PC. White noise (75 dB) was presented throughout habituation and testing to mask external noise. Before testing, the mice were habituated to both the injection procedure and the open-field arena for 30 min per day on three consecutive days. During each habituation session, each mouse received an intraperitoneal injection of saline and was habituated individually in the open-field arena for 30 min, with only one mouse in the arena at a time. Testing was conducted on two consecutive days. On day 1, both members of each pair received an intraperitoneal injection of either saline or L-368,899 at a dose of 10 mg/kg, according to their assigned treatment condition. On day 2, all mice received an intraperitoneal injection of saline. This fixed schedule was used to assess whether behavior on the day following L-368,899 administration approached that of the saline-treated control group. The control group received saline on both days, providing a comparison under the same repeated-testing schedule. On both test days, the mice were returned to their home cage immediately after injection and were placed together in the open-field arena 30 min later. Each test session lasted 30 min. Contact duration and mean inter-individual distance were quantified for each pair as measures of dyadic social interaction. Dominance relationships within the pairs were not assessed.

**Figure 5.**
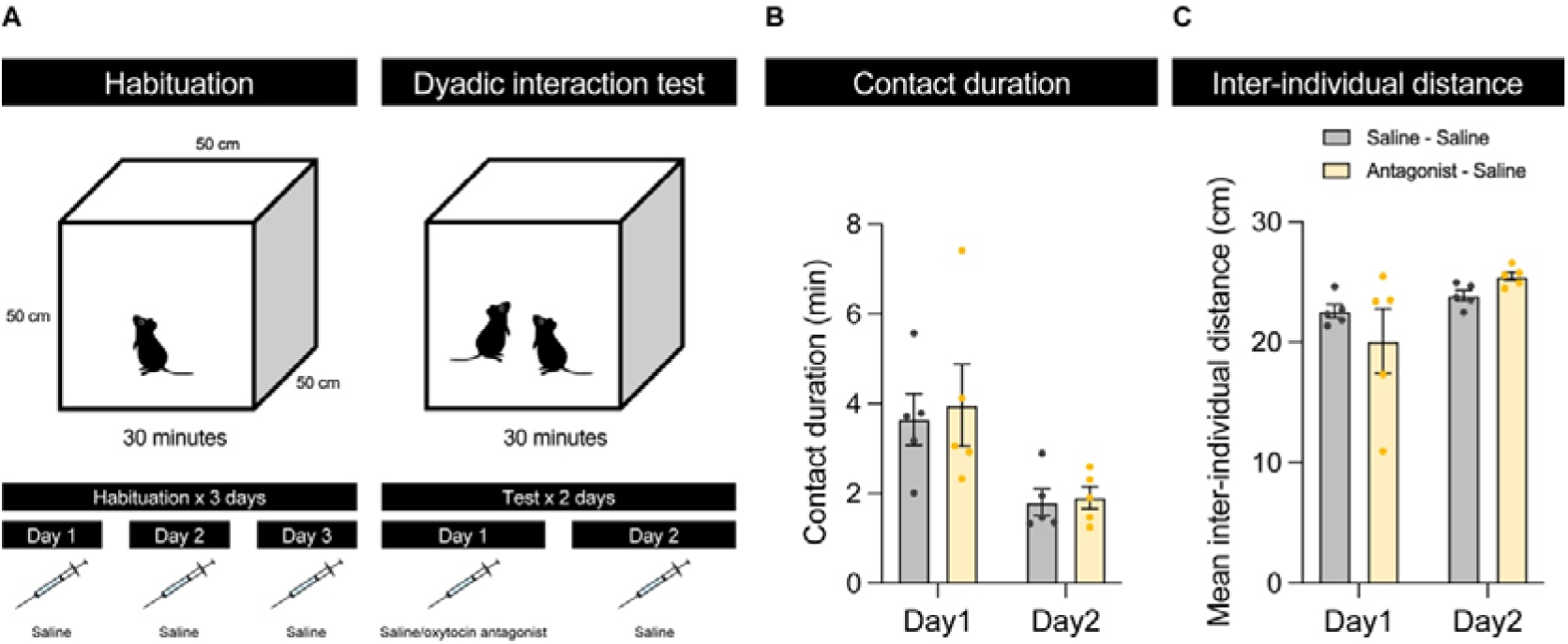
Effects of L-368,899 on contact duration and inter-individual distance during dyadic interactions between familiar male cage mates. A. Experimental procedure. Before testing, each mouse received saline and was habituated individually to the open-field arena for 30 min per day on three consecutive days. During testing, the two familiar cage mates were placed together in the arena. On Day 1, both members of each pair received an intraperitoneal injection of saline or L-368,899 (10 mg/kg) 30 min before testing. On Day 2, all mice received saline 30 min before testing. Each test session lasted 30 min. B. Contact duration between the two mice. Two-way mixed-design ANOVA detected a significant main effect of test day, with shorter contact duration on Day 2, but no significant main effect of treatment sequence or treatment-sequence-by-day interaction. C. Mean inter-individual distance. A significant main effect of test day was detected, with greater distance on Day 2, but neither the main effect of treatment sequence nor its interaction with day was significant. Data are presented as the mean ± SEM, with individual points representing pairs. Saline–saline: *n* = 5 pairs; L-368,899–saline: *n* = 5 pairs. The same pairs were tested on both days.

### 2.3. Drug

The oxytocin receptor antagonist L-368,899 hydrochloride (CAS No. 160312-62-9; Tocris Bioscience) was dissolved in saline, divided into aliquots, and stored at −80°C until use. Before each experiment, an aliquot was thawed, and the solution was administered intraperitoneally. L-368,899 was administered at a dose of 10 mg/kg in Experiments 1, 3, and 4 and at doses of 3 or 10 mg/kg in Experiment 2. Control mice received an equivalent volume of saline via the same route of administration. All injections were administered at a volume of 10 mL/kg body weight across all four experiments. The dose of 10 mg/kg was selected based on a previous study reporting reduced observational fear in male mice following intraperitoneal administration of L-368,899 30 min before testing (Pisansky et al., 2017). In Experiment 2, the additional 3 mg/kg condition was included to facilitate comparison with a previous study of sex preference using intraperitoneal L-368,899 administration in adult male mice (Haskal de la Zerda et al., 2020), rather than to test a specific prediction about differences between the two doses. Experiment 1 was restricted to 10 mg/kg because establishing and reassessing stable hierarchies required extensive testing, limiting the feasibility of additional dose conditions. In Experiment 3, no significant treatment effect was detected at 10 mg/kg, and testing was not extended to a lower dose. Experiment 4 was conducted as a complementary assessment of interactions between familiar cage mates and was limited to the 10 mg/kg condition rather than a dose–response evaluation. These dosing conditions were informed by previous behavioral studies in mice; brain exposure and oxytocin receptor occupancy were not measured in the present study.

### 2.4. Behavioral tracking and quantification

Behavioral data were processed using R via RStudio and GraphPad Prism. In Experiment 1, the numbers of wins and losses and the duration of each match were recorded manually by an observer blinded to treatment condition. For the analysis of rank outcomes, each cage was classified according to whether the treated mouse showed a change in numerical rank or became tied with another cage mate at any time from the administration day (Day 0) through Day 3, relative to its stable pretreatment position. Changes involving only untreated cage mates were not counted for this outcome. Because the same eight cages underwent both the first-ranked- and second-ranked-mouse treatment conditions, the occurrence of this outcome was compared between conditions using a two-sided exact McNemar test, with cage as the paired unit.

For the analysis of body weight in Experiment 1, the three measurements recorded for each mouse in the pretreatment period were averaged, as were the measurements recorded on the three days following L-368,899 administration (Days 1–3, with administration designated as Day 0). These individual means were compared using two-sided paired t-tests, separately for the first-ranked- and second-ranked-mouse treatment conditions (*n* = 8 mice per condition). Each mouse contributed one paired observation to the analysis. In Experiments 2–4, mouse positions were tracked from video recordings using Bonsai, an open-source visual programming framework for real-time and offline data processing (Lopes et al., 2015). Video images were converted to grayscale, smoothed, and inverted to generate monochrome images. Mice were detected by applying a contrast threshold, and their positions were estimated from the center coordinates of the detected body area. Locomotor activity was not analyzed. In Experiments 2 and 3, time spent in each stimulus zone was quantified as the cumulative time during which the center coordinate of the mouse’s detected body area was located within the 7.5-cm-wide zone adjacent to the corresponding stimulus basket. This measure represented zone occupancy rather than direct stimulus investigation. No additional criteria based on nose-to-stimulus proximity, head orientation, or active sniffing were applied. Consequently, facing away from the basket, grooming, remaining immobile, and passing through the zone were included whenever the positional criterion was met. In Experiment 4, the distance between the two mice was calculated from their tracked center coordinates. The mice were operationally classified as being in contact when the inter-individual distance was 5 cm or less. Contact duration was calculated as the cumulative time during which this criterion was met, and mean inter-individual distance was calculated across the entire 30-min test session.

## 3. Results

Experiment 1 examined rank trajectories following L-368,899 administration to first- or second-ranked mice in eight cages containing four mice each. Figure 1 summarizes the tube-test procedure and behavioral characteristics before antagonist administration. All cages established a linear hierarchy, with ranks remaining unchanged for three consecutive days before each treatment condition. The mean time required to meet this criterion was 12.6 days (SEM = 2.79) before the first-ranked-mouse condition and 7.1 days (SEM = 2.20) before the second-ranked-mouse condition. During the stable pretreatment period, matches between first- and second-ranked mice had the longest mean duration, whereas matches involving fourth-ranked mice were shorter (Figure 1F). Some matches involving the 1–3, 3–4, 2–4, and 1–4 rank pairings ended without direct physical contact (Figure 1G). Body weight was not significantly associated with social rank (Figure 1H, *R²* = 0.0017, *p* = 0.6903, simple linear regression; 96 measurements from 32 mice). These measurements characterize pretreatment performance and do not assess behavioral changes caused by L-368,899. Rank trajectories following antagonist administration are presented in Figure 2.

Following L-368,899 administration to first-ranked mice, the treated animals retained their first-ranked position without becoming tied with another cage mate in all eight cages, although rank changes occurred among some untreated cage mates (Figure 2A). Following administration to second-ranked mice, the treated animals showed heterogeneous trajectories (Figure 2B). In Groups #2 and #5, the treated mouse reached first rank on the administration day; it returned to second rank on the following day in Group #2 but remained at first rank throughout the remaining observation period in Group #5. In Group #1, the treated mouse fell to third rank on Day 1 and returned to second rank on Day 2. In Groups #6 and #7, the treated mouse became tied with the initially third-ranked cage mate on Day 3 and Day 1, respectively, with each mouse winning one of their two encounters. In Groups #3, #4, and #8, the treated mouse remained at second rank without becoming tied with another cage mate throughout the observation period. Changes in the treated animal’s numerical rank or the occurrence of a tie were observed in five of the eight cages following administration to second-ranked mice and in none following administration to first-ranked mice. There was a trend toward a higher frequency of these outcomes following administration to second-ranked mice, although the difference did not reach the conventional threshold for statistical significance (two-sided exact McNemar test, *p* = 0.0625).

Mean body weight during Days 1–3 following L-368,899 administration did not differ significantly from the pretreatment mean in either the first-ranked-mouse condition (*t*(7) = 0.258, *p* = 0.804) or the second-ranked-mouse condition (*t*(7) = 0.244, *p* = 0.814; two-sided paired t-tests; Supplementary Figure 1A–B). These comparisons used the mean of three measurements per period for each mouse, with eight mice in each condition.

Experiment 2 examined sex preference in male mice (Figure 3). A two-way mixed-design ANOVA revealed a significant main effect of stimulus (*F*(1, 21) = 10.60, *p* = 0.0038), but neither the main effect of drug (*F*(2, 21) = 2.83, *p* = 0.0814) nor the drug-by-stimulus interaction (*F*(2, 21) = 1.57, *p* = 0.2316) was statistically significant. After Šídák correction across the three within-group stimulus comparisons, saline-treated mice spent significantly more time in the female stimulus zone than in the male stimulus zone (adjusted *p* = 0.0112). A significant difference between stimulus zones was not detected in mice treated with either 3 mg/kg (adjusted *p* = 0.3442) or 10 mg/kg L-368,899 (adjusted *p* = 0.8118). Supplementary analyses likewise did not detect a treatment-group difference in female-minus-male zone occupancy (Welch test, *p* = 0.2909; permutation test, *p* = 0.2346). However, the within-group preference in saline-treated mice did not reach statistical significance in an exact sign-flip sensitivity analysis with Holm correction (adjusted *p* = 0.0703), indicating that this finding was sensitive to the analytical method. The present data therefore do not establish that L-368,899 reduced sex preference.

Experiment 3 examined preference for an unfamiliar male conspecific over a nonsocial object (Figure 4). A two-way mixed-design ANOVA revealed a significant main effect of stimulus (*F*(1, 16) = 53.71, *p* < 0.0001), whereas neither the main effect of drug (*F*(1, 16) = 0.145, *p* = 0.7084) nor the drug-by-stimulus interaction (*F*(1, 16) = 0.054, *p* = 0.8187) was statistically significant. After Šídák correction across the two within-group stimulus comparisons, mice spent significantly more time in the unfamiliar male stimulus zone than in the nonsocial object zone in both the saline-treated group (adjusted *p* = 0.000131) and the L-368,899-treated group (adjusted *p* = 0.000253). Thus, no statistically significant treatment-related difference in proximity-based social preference was detected under the present experimental conditions.

Experiment 4 examined whether systemic oxytocin receptor blockade alters dyadic interactions between familiar male cage mates in an open-field arena (Figure 5A; Supplementary Movie 4). Both members of each pair received an intraperitoneal injection of saline or L-368,899 (10 mg/kg) 30 min before testing on day 1. On day 2, all mice received saline 30 min before testing. Contact duration and mean inter-individual distance were measured during each 30-min session (Figure 5B and C). No statistically significant main effect of drug was detected for either contact duration (*F*(1, 8) = 0.1425, *p* = 0.7156) or inter-individual distance (*F*(1, 8) = 0.0888, *p* = 0.7732). A significant main effect of experimental day was detected for both measures: contact duration decreased from day 1 to day 2 (*F*(1, 8) = 11.21, *p* = 0.0101), whereas inter-individual distance increased (*F*(1, 8) = 6.216, *p* = 0.0373). The experimental day × drug interaction was not statistically significant for either contact duration (*F*(1, 8) = 0.03277, *p* = 0.8609) or inter-individual distance (*F*(1, 8) = 2.332, *p* = 0.1653). Thus, no statistically significant effect of L-368,899 on either measure of dyadic interaction was detected under the present experimental conditions.

## 4. Discussion

In the present study, we examined the effects of systemic administration of the oxytocin receptor antagonist L-368,899 across multiple domains of social behavior in male mice. The antagonist did not produce a uniform reduction in social approach. Instead, different behavioral patterns were observed across tasks and, in the tube test, according to the hierarchical position of the treated animal. Administration to first-ranked mice was not followed by changes in their rank, whereas rank fluctuations were observed more frequently after administration to second-ranked mice. However, the difference in the frequency of rank changes between these conditions did not reach statistical significance and should therefore be interpreted as preliminary evidence of a possible rank-dependent response. In the sex-preference task, preference for the female stimulus zone over the male stimulus zone was statistically detectable in saline-treated subjects but not in L-368,899-treated subjects; the stimulus mice received no injections. Nevertheless, the stimulus-by-drug interaction was not significant, precluding a definitive conclusion that the antagonist reduced sex preference. No statistically significant drug effects were detected in either general social preference, in which only the subject mouse was treated, or dyadic interactions between familiar male cage mates, in which both animals received the same treatment. These findings motivate further investigation of possible task- and rank-related differences in responses to L-368,899, but do not establish that social context itself accounts for the patterns observed across tasks.

In the tube test, first-ranked mice retained their position after L-368,899 administration in all eight cages examined. Rank changes were more frequently observed when the antagonist was administered to second-ranked mice, although the comparison between the first- and second-rank conditions did not meet the conventional threshold for statistical significance. Body weight was unchanged following antagonist administration, providing no evidence that the observed rank fluctuations were attributable to an acute deterioration in general physical condition. In addition, all animals underwent comparable handling and injection procedures: the target-ranked mouse received L-368,899, whereas the remaining cage mates received saline. Nonspecific injection stress alone is therefore unlikely to explain why rank fluctuations were observed more often following treatment of second-ranked animals. Although all cages met the same pretreatment criterion of stable ranks for three consecutive days, we did not establish whether differences in baseline competitive relationships or individual behavioral characteristics were associated with subsequent rank trajectories. Furthermore, the first-ranked-mouse condition always preceded the second-ranked-mouse condition, preventing separation of rank-related differences from effects of testing order or prior experience. Given these limitations, the small number of cages, and the inconclusive statistical evidence, the present data do not establish that oxytocin receptor blockade destabilizes intermediate social rank.

If the apparent rank-dependent pattern is replicated, one possible explanation is that mice occupying different hierarchical positions differ in their baseline oxytocin receptor signaling. Oxytocin receptor densities in several brain regions vary according to social status (Lee et al., 2019), raising the possibility that the same systemic receptor blockade produces different behavioral consequences in first- and second-ranked mice. An alternative, but not mutually exclusive, explanation concerns the distinct behavioral demands associated with different hierarchical positions. First-ranked mice interact exclusively with subordinate cage mates, whereas second-ranked mice encounter both a dominant individual and lower-ranked individuals. Maintaining an intermediate position may therefore require greater adjustment of behavior according to the identity and rank of the opponent. Under this interpretation, oxytocin receptor blockade could affect the processing or use of socially relevant information without directly altering dominance motivation or physical competitive ability. This explanation remains hypothetical because individual recognition, opponent-specific behavioral selection, and aggressive or affiliative behaviors during individual matches were not quantitatively assessed.

The sex-preference findings also require cautious interpretation. In the ANOVA-based comparisons, saline-treated males spent significantly more time in the female stimulus zone than in the male stimulus zone, whereas a statistically significant within-group difference was not detected after treatment with either dose of L-368,899. The stimulus-by-drug interaction was not statistically significant. Thus, the presence of a statistically significant preference in one treatment group and its absence in another does not by itself demonstrate a significant pharmacological reduction in preference. Moreover, the within-group preference in saline-treated mice did not reach statistical significance in the Holm-adjusted sign-flip sensitivity analysis, indicating that this finding depended on the analytical method. The descriptive pattern is broadly consistent with previous pharmacological findings obtained using L-368,899 (Haskal de la Zerda et al., 2020) and with altered social behavior reported in oxytocin receptor-deficient mice (Sala et al., 2013; Takayanagi et al., 2005). Studies of oxytocin-deficient mice, however, have produced different patterns of social approach and recognition (Donaldson and Young, 2008; Choleris et al., 2003; Crawley et al., 2007). Oxytocin receptor signaling could contribute to several processes relevant to performance in this task, including sexual motivation, sex discrimination, and the assignment of motivational value to opposite-sex social cues (Boccia et al., 2007; Haskal de la Zerda et al., 2020; Carter, 1992; Oti et al., 2021). The present experiment was not designed to distinguish among these possibilities, and a larger confirmatory study will be necessary to determine whether L-368,899 reliably alters sex preference.

No statistically significant effects of L-368,899 were detected in either the social-preference task or the direct dyadic interaction test between familiar male cage mates. In the social-preference task, only subject mice received injections, whereas stimulus mice received no injections, and mice spent significantly more time in the zone adjacent to the unfamiliar male conspecific than in the zone adjacent to the nonsocial object under both treatment conditions. In the dyadic interaction test, both members of each pair received the same treatment, and neither contact duration nor mean inter-individual distance differed significantly between saline- and L-368,899-treated pairs. In line with previous studies reporting that disruption of oxytocin signaling affects some, but not all, measures of social approach and recognition (Ferguson et al., 2000; Haskal de la Zerda et al., 2020; Choleris et al., 2003; Crawley et al., 2007), these findings suggest that oxytocin receptor signaling does not uniformly regulate all forms of social behavior. Contact duration decreased and mean inter-individual distance increased from the first to the second test day in both treatment groups. These changes may reflect habituation to, or prior experience with, the testing environment. The fixed treatment schedule was intended to assess whether behavior on the day following L-368,899 administration approached that of the saline-treated controls. Although the saline–saline group provided a comparison for changes associated with repeated testing, the design was not a counterbalanced crossover, and the interval between sessions was not established as an adequate pharmacological washout. Residual drug effects or lasting consequences of the initial treatment therefore cannot be excluded on day 2. Moreover, because no significant treatment effect was detected, the day-2 observations should not be interpreted as demonstrating recovery from a drug-induced behavioral change.

The absence of statistically significant drug effects in these two tasks should not, however, be interpreted as definitive evidence that oxytocin receptor signaling is unnecessary for general affiliative behavior. Conventional null-hypothesis tests do not establish equivalence between treatment conditions, particularly with modest sample sizes. Rather, the present findings indicate that any effects of acute systemic L-368,899 on these measures were not sufficiently large or consistent to be detected under the current experimental conditions. Reporting effect sizes and confidence intervals, and applying equivalence or Bayesian analyses in future work, would help determine the range of drug effects compatible with these data.

One framework that may account for the different behavioral patterns across tasks is that oxytocin receptor signaling contributes more strongly when behavior requires the discrimination and context-dependent use of specific social information. Second-ranked mice may need to adjust their responses according to whether an opponent is dominant or subordinate, while sex preference requires discrimination between male and female conspecifics. By comparison, preference for a conspecific over an object and interaction with a familiar cage mate may place different demands on social discrimination and behavioral flexibility. However, the present experiments differed not only in social context but also in the behaviors measured, task procedures, and treatment configurations. Differences in the sensitivity of the outcome measures may also contribute to the observed patterns. The cross-task comparisons therefore cannot isolate social context as the source of variation in drug responses. A more direct test would measure the same behavior under systematically varied social conditions—for example, assessing the same measures of sexual behavior with familiar versus unfamiliar partners while keeping drug administration and testing procedures constant. The proposed framework should thus be regarded as a hypothesis for future testing rather than a mechanism demonstrated by the present experiments.

Chemosensory processing represents one possible pathway through which oxytocin receptor signaling could influence context-dependent social behavior. Rodents rely heavily on olfactory and other chemosensory cues to discriminate among individuals and between sexes. Oxytocin receptors are expressed in brain regions involved in olfactory processing (Vaccari et al., 1998), and pharmacological inhibition of oxytocin signaling in the anterior olfactory nucleus impairs social odor recognition without disrupting nonsocial odor discrimination (Oettl et al., 2016). Mice can also infer the social status of unfamiliar individuals from chemosensory information during the tube test (Borak et al., 2025). Systemic oxytocin receptor blockade could therefore alter either the processing of social odors or the behavioral use of chemosensory information. However, because olfactory detection, social odor discrimination, and individual recognition were not directly assessed, the present findings cannot distinguish an impairment in chemosensory processing from changes in social motivation or response selection.

Oxytocin signaling has also been implicated in social recognition and social memory (Ferguson et al., 2000; Lukas et al., 2013). The absence of a statistically significant effect on interactions between familiar cage mates argues against a severe, generalized disruption of social engagement with familiar conspecifics under the present experimental conditions. However, the familiar dyadic interaction test did not directly assess either individual recognition or social recognition memory. Future studies should therefore include explicit individual-recognition and social-memory paradigms to determine whether the possible effects of oxytocin receptor blockade on rank stability and sex preference arise from altered recognition, motivation, or behavioral selection.

The doses of L-368,899 used in this study were selected based on previous behavioral studies in mice. Intraperitoneal administration of 10 mg/kg reduced observational fear when administered 30 min before testing (Pisansky et al., 2017), and administration of 3 mg/kg has been associated with an altered pattern of sex preference in adult male mice (Haskal de la Zerda et al., 2020). Other studies provide further evidence of behavioral actions of this compound. In sexually experienced male rats, L-368,899 reduced time spent near a receptive female while leaving a significant female preference detectable in both treatment groups (Blitzer et al., 2017). In mice, L-368,899 attenuated the acquisition, but not retrieval, of lithium chloride-induced conditioned taste aversion (Olszewski et al., 2013). These findings support the behavioral activity of L-368,899 across experimental settings, but do not establish equivalent receptor blockade across species, doses, or tasks. Peripherally administered L-368,899 also enters the cerebrospinal fluid and accumulates in several brain regions in rhesus monkeys (Boccia et al., 2007), supporting the possibility of central actions following systemic administration. However, brain exposure and receptor occupancy were not measured in the present mice, leaving the extent and duration of receptor blockade during each task uncertain. Systemic administration also cannot identify the neural sites responsible for the observed patterns or exclude contributions from peripheral oxytocin receptors. Accordingly, a nonsignificant effect in a particular task or rank group should not be interpreted as evidence that oxytocin receptor signaling is unnecessary for that behavior.

Several limitations should be considered. First, the numbers of cages and animals were modest, and the study may have had insufficient statistical power to detect treatment effects or interactions between treatment and either rank or stimulus. The apparent rank-dependent effect and the descriptive pattern observed in the sex-preference task should therefore be regarded as hypothesis-generating findings that require independent replication. Second, we examined drug administration only in first- and second-ranked mice in Experiment 1. Whether oxytocin receptor blockade produces similar or different effects when administered to third- or fourth-ranked individuals remains unknown. Furthermore, dominance relationships were not assessed within the pairs in Experiment 4, preventing evaluation of whether rank influenced responses during dyadic interactions. Third, we examined only the effects of acute systemic administration of a single oxytocin receptor antagonist, L-368,899. This approach cannot distinguish central from peripheral effects, identify the relevant brain regions and neural circuits, or exclude compound-specific effects, including potential off-target actions. The present findings therefore do not establish a general role for the oxytocin system across the behaviors examined. Studies using structurally distinct oxytocin receptor antagonists and receptor-specific genetic approaches will be needed to determine whether the observed patterns reflect oxytocin receptor inhibition. Local pharmacological manipulations, oxytocin receptor agonists, and repeated treatment could further clarify the sites of action and the effects of different forms of oxytocin receptor modulation. Fourth, only Experiment 2 included both 3 and 10 mg/kg conditions; Experiments 1, 3, and 4 were limited to 10 mg/kg. The dose–response relationships therefore remain incompletely characterized, and the absence of significant effects at 10 mg/kg does not exclude effects at lower doses. Fifth, Experiments 2 and 3 measured time spent in predefined stimulus zones rather than direct stimulus investigation. This measure captures spatial preference but does not distinguish active stimulus-directed investigation from other behaviors occurring within the same zone. These findings should therefore be interpreted in terms of proximity-based preference. Future studies combining zone occupancy with measures of nose-to-stimulus proximity, head orientation, and active sniffing could clarify whether these measures reveal similar or distinct effects of L-368,899. Sixth, Experiment 4 used a fixed treatment schedule to examine whether behavior on the following day approached that of the control group. Because treatment order was not counterbalanced and an adequate washout period was not established, order, habituation, and carryover effects cannot be excluded. Moreover, the absence of a significant treatment effect on Day 1 precludes interpreting the Day 2 findings as evidence of recovery from a drug-induced change. Seventh, we did not directly assess olfactory function, individual recognition, social memory, sexual motivation, or opponent-specific behavioral strategies. Alterations in these processes therefore remain possible explanations for the observed behavioral patterns rather than demonstrated mediators. Eighth, the estrous cycle of the female stimulus mice was neither monitored nor experimentally controlled and may have contributed to variability in male responses to the female stimuli. Finally, only male subjects were examined. Whether comparable context- and rank-dependent patterns occur in females remains unknown, although female mice can also form stable social hierarchies (Fulenwider et al., 2024; LeClair et al., 2021).

In conclusion, systemic administration of L-368,899 was associated with different behavioral patterns across tasks assessing social rank, sex preference, general social preference, and familiar dyadic interactions in male mice. First-ranked mice maintained their hierarchical positions following treatment, whereas rank fluctuations were observed more frequently when second-ranked mice were treated; however, the difference between the two rank conditions was not statistically conclusive. Similarly, a statistically significant preference for the female stimulus zone was detected in saline-treated subjects but not in L-368,899-treated subjects, although the stimulus-by-drug interaction was not statistically significant. No statistically significant effects were detected on proximity-based preference for an unfamiliar male conspecific over a nonsocial object or on dyadic interactions between familiar male cage mates. These findings provide no evidence of a generalized suppression of social approach under the conditions examined and identify possible task- and rank-related patterns that require independent replication. Because the tasks differed in both social context and behavioral measures, the contribution of context itself remains unresolved. Future experiments measuring the same behavior across systematically varied social contexts will be needed to test whether responses to L-368,899 are context-dependent.

## Data availability

The data supporting the findings of this study are available from the corresponding author upon reasonable request.

## Code availability

The original codes written for the analyses are available from the corresponding author upon reasonable request.

## Acknowledgements

This research was supported by JSPS KAKENHI 18KK0070 (KT), 19H05316 (KT), 19K03385 (KT), 19H01769 (KT), 20J21568 (KY), 22H01105 (KT), 23H02787 (KT), 23K27478 (KT), 23K22376 (KT), 24H00729 (KT), 24K16869 (KY), 24KJ0069 (KY), 24K06626 (KH), and 25KJ0306 (KH), Keio Gijuku Fukuzawa Memorial Fund for the Advancement of Education and Research (KT), HOKUTO Foundation for the Promotion of Biological Science, and Smoking Research Foundation (KT). We thank Kohei Yamamoto, Shohei Kaneko, Yasuyuki Niki, Mizuki Yamamoto, Haruki Kasahara, and Hiroto Inoue for their assistance and valuable discussions. We also thank Dr. Takaaki Ozawa and Takahide Omori for their valuable comments on the manuscript.

## Author contributions

DN and KT designed all the experiments. DN and SM collected the tube test data. DN and KT collected data on sex preference, social preference, and dyadic interaction tasks with the help of HH, SY, and KH. HH, RT, and YU helped to set up the experimental apparatus. DN, SM, and KT analyzed the data. KT wrote the manuscript. DN and KT created all the figures. DN, SM, KY, YU, KH, and KT discussed the data and commented on the manuscript accordingly. KT revised the manuscript.

